# Context-dependent dynamics lead to the assembly of functionally distinct pitcher-plant microbiomes

**DOI:** 10.1101/727701

**Authors:** Leonora S. Bittleston, Matti Gralka, Gabriel E. Leventhal, Itzhak Mizrahi, Otto X. Cordero

## Abstract

Niche construction through interspecific interactions can condition future community states on past ones. However, the extent to which such history dependency can steer communities towards functionally different states remains a subject of active debate. Using bacterial communities collected from wild pitchers of the carnivorous pitcher plant, *Sarracenia purpurea*, we tested the effects of history on composition and function across communities assembled in synthetic pitcher plant microcosms. We found that the diversity of assembled communities was determined by the diversity of the system at early, pre-assembly stages. Species composition was also contingent on early community states, not only because of differences in the species pool, but also because the same species had different dynamics in different community contexts. Importantly, compositional differences were proportional to differences in function, as profiles of resource use were strongly correlated with composition, despite convergence in respiration rates. Early differences in community structure can thus propagate to mature communities, conditioning their functional repertoire.

## Introduction

Microbes profoundly shape our ecosystems, yet we still lack a clear understanding of the processes driving community assembly and related ecosystem functioning^1^. Community assembly is difficult to predict because the dynamics of any particular species can be dependent on the community context; niches are created or destroyed through biotic interactions with other members of an assembling community^2–4^. Microbial communities are complex, with many species and different kinds of interactions among species. For example, microbes can facilitate other species’ growth via excretion of metabolic waste products^5–8^, or actively interfere with their growth through the production antimicrobial compounds^9^. Microbes can also engage in strong cooperative interactions, whereby energy transducing metabolic interactions are coupled across species^10^. This diversity of interactions creates many potential contexts for species dynamics, implying that the behavior of one species is dependent on the background of interactions. Stochastic changes in the biotic context—for example, priority effects and random colonization or extinction events—can have long-lasting consequences for community structure^3^. As a result, microbial communities can reach different compositional states due to variation in biotic context in addition to variation in abiotic environmental conditions such as weather events or resource pulses. These history-dependent effects are collectively called “historical contingencies”^2–4^.

The extent to which historical contingency leads to alternative community states^2,11^ remains an active subject of debate. In a strongly selective environment historical contingencies may not matter such that communities converge to the same compositional and functional outcomes^12^. For example, bacterial communities from widely different sources converged reproducibly in single-carbon source synthetic media^7^. Beyond microbes, Mediterranean plant communities in environments with frequent fires were formed by related groups of species that share key traits^13^. In contrast, other studies have found that historical contingency leads to different community compositions and functions, for example: priority effects led to large differences in ecosystem function for wood-decaying fungi^14^ and productivity of grassland plants^15^. A third, and perhaps largest, set of studies has found convergence in terms of function but not species (or phylogenetic) composition, for example: grassland plants^16^, the stratified layers of a hypersaline microbial mat^17^, microbial communities colonizing the surface of seaweed^18^, and the bacteria and archaea living in bromeliad tanks^19^.

Functional convergence without species convergence is more likely when the functions being measured are performed by many species from different lineages, and thus are redundant within the broader species pool. Here, again, there are contrasting reports in the literature on the prevalence of functional redundancy. A number of studies have found functional redundancy in microbial communities^19,20^, while others emphasize important functional differences that depend on species composition^21^. The degree of redundancy is clearly related to the function and system examined; for example, aerobic respiration is found across many bacteria, while the ability to degrade lignin is rare. Thus, ‘narrow’ functions, such as the hydrolysis of complex carbon compounds, are generally carried out by rare community members^22^. When relevant functions are variable across genetic backgrounds (not highly redundant) and dependent on interactions, historical contingencies could have major effects on the functional capabilities of a community and on nutrient cycling within ecosystems^14^. Therefore, studies in microbial ecology need to address more specific, relevant functional measurements and examine not just on historically contingent *compositional* states, but also historically contingent *functional* states.

The modified leaves of carnivorous pitcher plants host small ecosystems composed of bacteria, fungi, protozoa, rotifers and arthropods^23,24^ and present an excellent system to investigate the role of historical contingency on function and composition. Bacteria, in particular, are thought to assist their pitcher plant hosts in breaking down captured prey^25–28^, creating a clear link between a relevant ecosystem function, the degradation of insect material, and the microbial species composition. Using bacterial communities from ten wild pitchers of the purple pitcher plant, *Sarracenia purpurea*, we tested to what extent historical contingencies would impact community assembly and substrate degradation. To this end, we transferred the communities from living pitcher plants into *in vitro* microcosms and performed a community assembly experiment, whereby communities are serially passaged until a stable composition is reached. During this assembly process, microcosms can converge or remain distinct due to differences in the initial species pool as well as to differences in biotic interactions across the microcosms. By comparing the assembly dynamics of ten different plant microbiomes we asked to what extent communities converge to a single compositional and functional state, and whether any lack of convergence could be explained by historical contingency.

## Results

### Distinct stable communities assemble in microcosms

The aquatic communities from 10 individual *Sarracenia purpurea* pitchers were filtered to focus on bacteria and inoculated into a realistic, complex nutrient source: sterilized, ground crickets in acidified water. The *in vitro* communities were serially transferred every three days for 21 transfers, using a low dilution rate of one-part culture to one-part fresh media. Community composition was measured for each transfer using 16S rRNA sequencing (see Methods). From ~8 million sequences across all microcosms and timepoints, DADA2 analysis inferred 889 distinct ASVs (Amplicon Sequence Variants, which we treat as units of diversity). The most abundant phylum was Proteobacteria, followed by Firmicutes and Bacteroidetes. The top twenty ASVs are displayed in Figure 1a, accounting for 69.4% of the reads; for ASVs without assigned genera, genus names were recovered from the full 16S rRNA genes of our cultured strains that matched 100% with ASV sequences. The most abundant genera (*Aquitalea*, *Pseudomonas*, *Achromobacter*, *Comamonas* and *Delftia*), all contain bacterial species known to live in freshwater, soil, or plant-associated habitats^29–33^. Using DNA concentrations as a proxy to measure biomass increase, we found that there is a ~10-fold increase in biomass during in-vitro assembly (Figure 1a). Microcosms M03 and M09 stood out as those with the highest biomass yield as well as having the lowest diversity.

**Figure 1.**
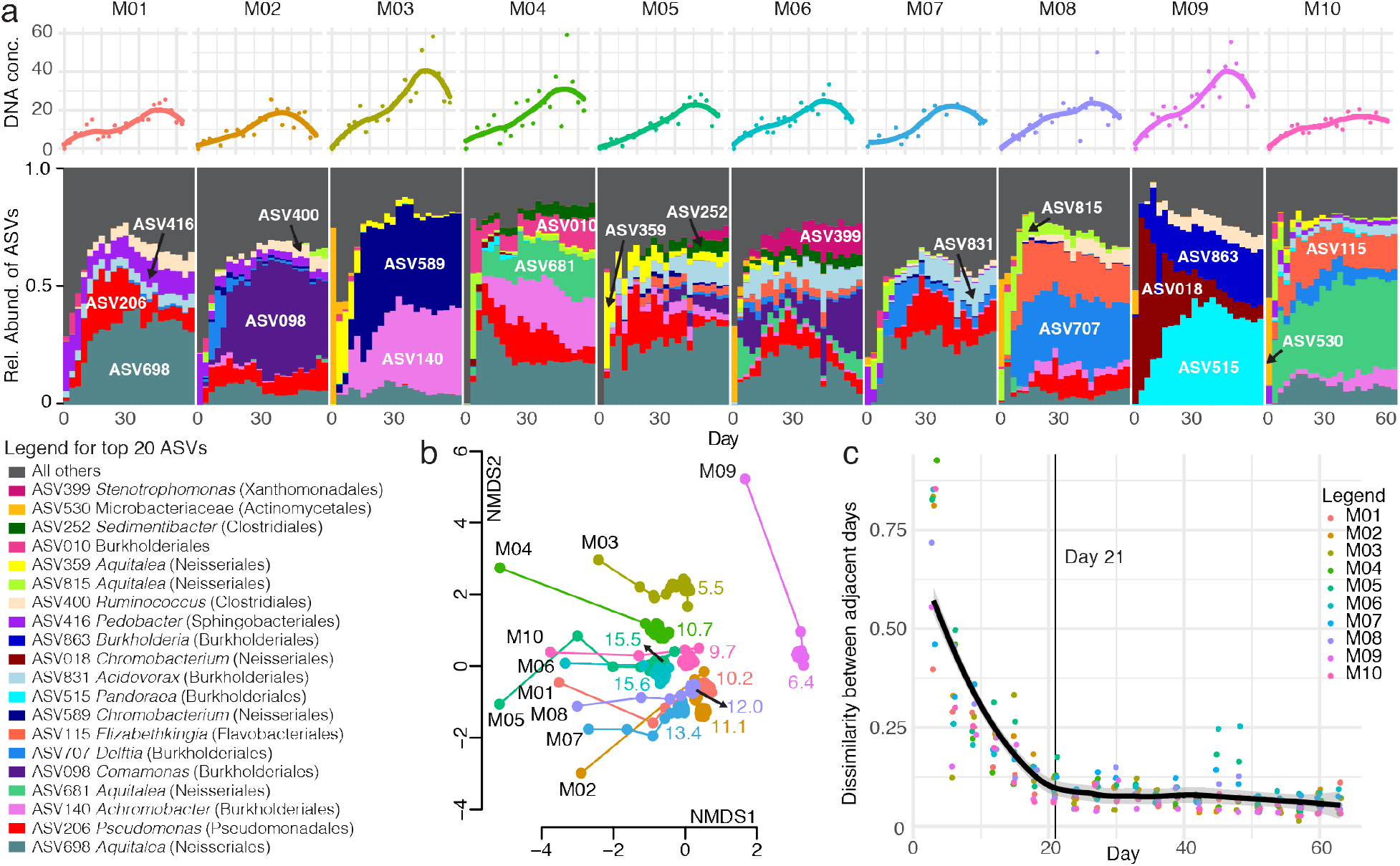
Microcosm communities approach distinct equilibria. a) Relative abundances of the top 20 ASVs across the ten microcosms during the course of the serial transfer experiment. ASVs are listed one time each on the bar plot, with taxonomic classification in the legend. The DNA concentrations for each timepoint are graphed above as points, with a Loess fit as a solid line. b) Communities change quickly and then stabilize in the Non-metric Multidimensional Scaling (NMDS) plot of Bray-Curtis dissimilarities of the microcosm community compositions. The microcosm name is listed in black next to the Day 0 community of each microcosm, and the lines connect the timepoints. Colored numbers indicate the mean effective number of species for each community post-Day 21. c) The Bray-Curtis dissimilarity of ASV relative abundances between adjacent days decreases over the course of the experiment. The thick black line shows a Loess fit to all data points, and the thin line marks Day 21.

The community assembly dynamics were dominated by both ASV loss and dramatic changes in ASV abundances. We assessed coarse-grained dynamics using the Bray-Curtis dissimilarity between subsequent time points, which exhibited two seemingly distinct phases (Figures 1b and 1c): a first phase of rapid change, where many species went extinct and the survivors increased in frequency; and a second phase of slow changes that started after 7 transfers (21 days, indicated by the thin line in Figure 1c), in which extinctions were rare and likely caused by competitive interactions. In this last phase, communities slowly approached what seemed to be an equilibrium state (Figure 1b and 1c). However, the near-equilibrated communities maintained significant differences both in terms of composition and diversity. For example, no microcosm had the same top three ASVs (Figure 1a), the NMDS ordination showed generally non-overlapping points for the different microcosms (Figure 1b), and, even after Day 21 the effective number of species (see Methods for calculation) still differed among the microcosm communities, ranging from about 6 to 16 (small colored numbers in Figure 1b). In summary, communities tended to first converge due to the fast loss of diversity (primarily along the first NMDS axis), but remain distinct due to historical contingencies (primarily along the second NMDS axis).

Interestingly, the differences in richness near-equilibrium were seemingly pre-determined at early stages of assembly. Communities lost many ASVs between days 0-3 during initial adjustment to the laboratory environment (Figure 2a), but the richness measured from the first timepoint of the experiment (Day 3) was an excellent predictor of the richness at the end of the experiment (Day 63) with R^2^ = 0.9008 and p < 0.0001 (Figure 2b). Notably, richness in samples taken directly from the pitcher plant (Day 0) had no significant correlation with that of Day 63 (R^2^ = 0.1978, p = 0.1105), consistent with the notion that some ASVs in the pitcher plant fluid that were either metabolically inactive or unable to grow in our experimental conditions. Thus, changes in community composition after only three days of adjusting to the lab environment propagated throughout the assembly process, suggesting that historical contingencies played a significant role in structuring communities.

**Figure 2.**
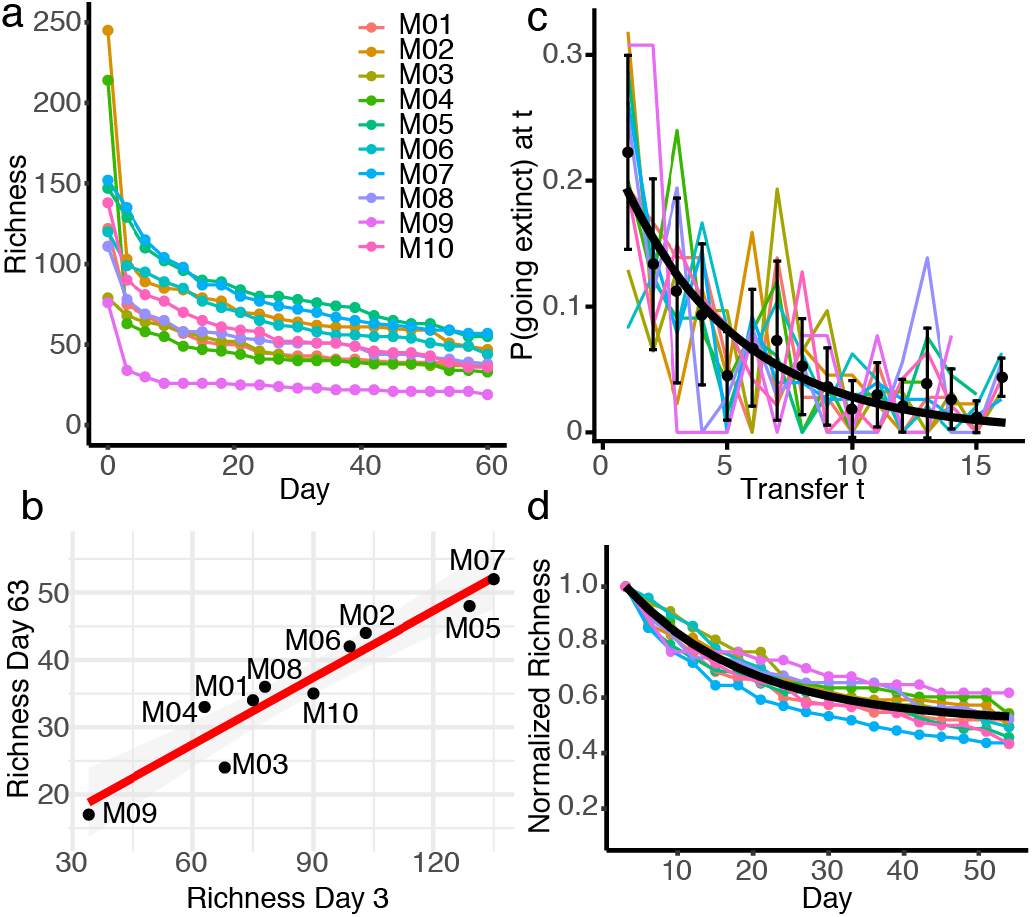
Early richness predicts final richness and communities equilibrate at a common rate. a) Richness over time for each microcosm community. b) Richness on Day 3 (1^st^ timepoint) is strongly correlated with richness on Day 63 (21^st^ timepoint). Linear model: R^2^ = 0.9008, p < 0.0001. c) The probability of going extinct at transfer *t*. Colored lines are probability densities for individual microcosms. Black points are averages across microcosms, the black line is the maximum likelihood distribution with a common parameter across microcosms (see main text and Methods) given by the inverse mean extinction time. d) Richness over time for each microcosm community, normalized by the richness on Day 3. The black line shows the exponential decay curve parametrized by the mean proportion of surviving ASVs (from panel b) and the common ASV extinction rate (from panel c).

Given the strong correlation between initial and final richness, we wondered whether temporal dynamics of community equilibration were also correlated across microcosms. For example, the temporal dynamics of ASV loss could be driven by external factors such as transfer intervals and the dilution factor, in which case all microcosms should exhibit comparable dynamics. Alternatively, the dynamics might be driven by community context and thus specific to each microcosm. To answer this question, we measured the distribution of ASV extinction times across the microcosm communities (Fig. 2c). As a null model, we tested if the loss of ASVs is a random process where each ASV that is bound to go extinct in a given community context does so with a fixed probability *p* in each transfer. This assumption implies that the extinction time distribution is described by a geometric distribution, which indeed gave a good fit for all microcosms (Pearson’s chi-squared test for differences was not significant: p > 0.05 in all cases). To investigate if microcosms can be described by a common extinction time distribution, we used the Akaike Information Criterion to compare two models: using either one parameter per microcosm or a single parameter describing all microcosms. Surprisingly, the single-parameter model was strongly favored (relative likelihood of ~1000) suggesting that temporal dynamics of ASV extinction in our communities are the same and likely driven by external factors rather than biotic context. This ‘universal’ dynamic of species loss is also revealed when studying relative richness, i.e. normalized by richness at Day 3 (Figure 2d). After this normalization, all relative richness curves mapped well to our model’s maximum likelihood distribution and approached a common equilibrium relative richness level (approximately ~50% of the initial richness, regardless of its absolute value, Figure 2d). Thus, about half of all initial ASVs were doomed to go extinct at a random point during the assembly process, while the rest persisted indefinitely.

We investigated if community context influenced the dynamics of individual ASVs. Although the ten microcosms reached different compositional states, on average ~90% of the communities were composed of ASVs that occurred in more than one microcosm (Supplementary Figure S1), and this overlap allowed us to ask to what extent the same species behave similarly or not when their community context changes. To this end, we first looked at the extinction times and found that the majority of the shared ASVs dropped out of different microcosms at different times (Figure 3a), often with large differences in their persistence time. For example, ASV681 (in Figure 3b) dropped out by Day 12 in microcosm M08, but persisted at high relative abundance through the end of the experiment in microcosms M04, M06 and M10. We focused on ASVs that persisted at least once past the 7^th^ transfer (Day 21) and found that although the same ASV can act differently in different community contexts (Figure 3b), the temporal trajectories of the same ASVs in different microcosms tended to be somewhat more correlated with each other than randomly chosen ASVs (Fig. 3c). This result implies that, as one might expect, species identity determines species abundance dynamics in different biotic contexts to some degree. However, only 55% of these ASVs had significantly correlated abundance trajectories in at least two microcosms and only 3% were significantly correlated across all their microcosms (Fig. 3d, see Methods for details). Furthermore, shared ASVs did not always have the same fate: about 80% of ASVs present in two microcosms either dropped out or persisted in in both, and about 50% of ASVs in 6 microcosms had the same fate in all cases (Fig. 3e). These results indicate that in most cases biotic context and interactions with other community members shaped ASV dynamics.

**Figure 3.**
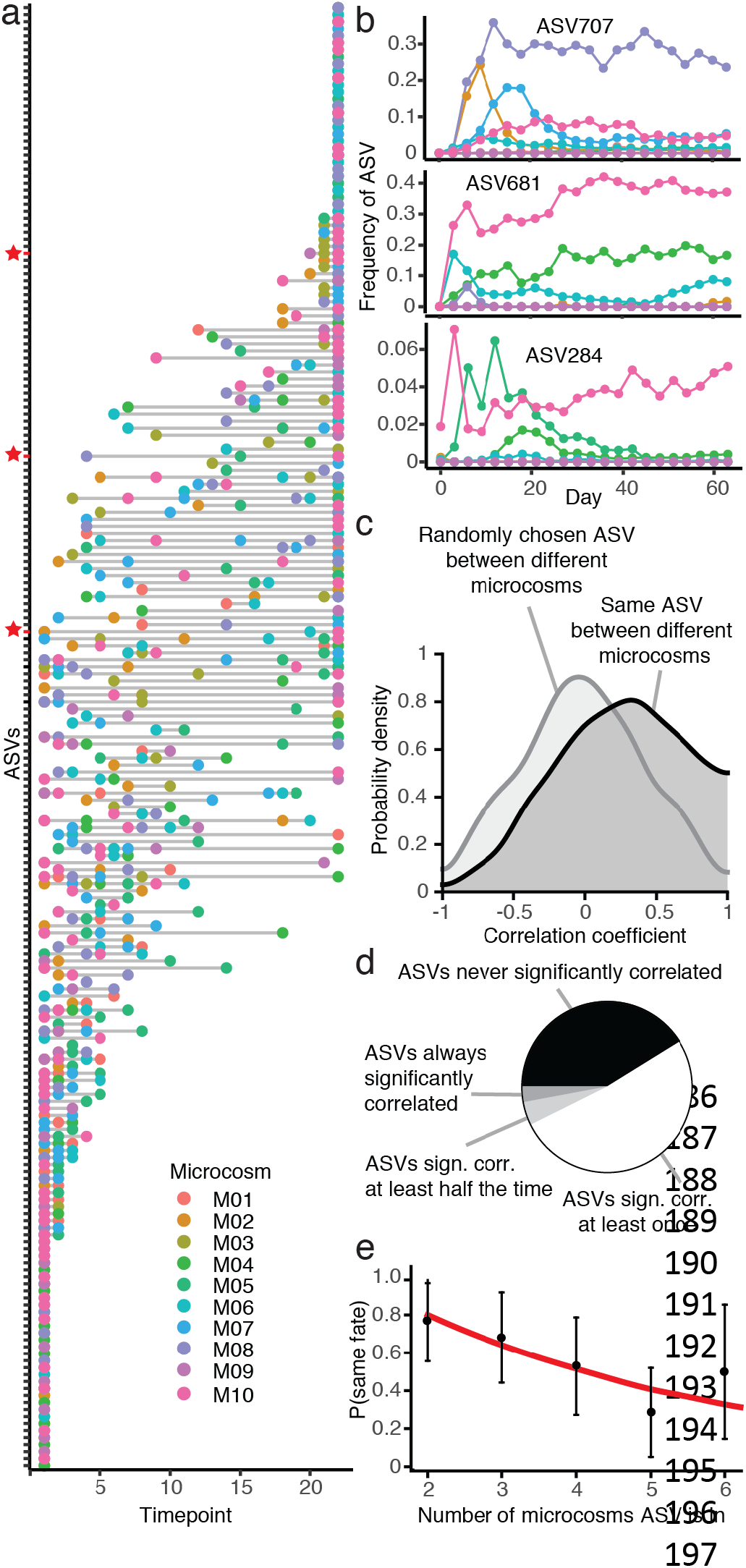
Species dynamics are context dependent. a) The y-axis lists all ASVs shared by at least two microcosms. Colored points mark which day the ASV was lost from that particular microcosm. Points at the far right of the graph are ASVs that persisted to the end of the serial transfer experiment. Red stars mark the ASVs shown in panel b. b) Three examples of ASV dynamics in the different microcosms over time, where ASVs were lost early in some microcosms but persisted until the end in others. c) Probability density of correlation coefficients for the correlation between the same ASVs in different microcosms versus randomly chosen ASVs. d) Pie chart showing proportions of ASVs that are never, at least once, at least half of the time or always significantly correlated in different microcosms. e) Probability of ASVs having the same fate (either persisting or going extinct) in different microcosms depending on the number of microcosms they are present in. Red line shows null expectation (see Methods).

We found that the effects of historical contingency on community assembly were remarkably consistent and reproducible. We performed the same experiment on a second set of microcosms from the same inocula that were not subjected to initial filtering, but otherwise underwent the same transfer protocol and amplicon sequencing. However, despite likely differences in protozoan predation (particularly for microcosms M02, M05 and M06 where observed activity persisted for at least 15 days), the unfiltered and filtered microcosm bacterial communities followed the same trajectories in community composition (see NMDS plot of Bray-Curtis dissimilarities, Supplementary Figure S2). The effects of initial pitcher source environment and composition persisted and were reproducible across both our filtered and unfiltered samples.

### ASV composition is highly correlated with substrate utilization

Functional redundancy is thought to be widespread in regional pools of bacteria^34^, and thus communities can have diverse compositions but converge to similar functional activity. Previous studies have generally found stronger convergence in function than in composition, and this was a likely outcome from our experiment as all microcosms experienced the same environment in terms of nutrients, temperature and light. In agreement with this, carbon dioxide production, as measured with the MicroResp system, was highly variable across the different microcosms for the first measurement (Day 0 – 3), but then quickly converged to a similar level (Figure 4a). After Day 3, communities could not be reliably distinguished based on their CO_2_ output. The low variation among microcosms in percent CO_2_ suggests that the bacterial communities were respiring at about the same rate.

**Figure 4.**
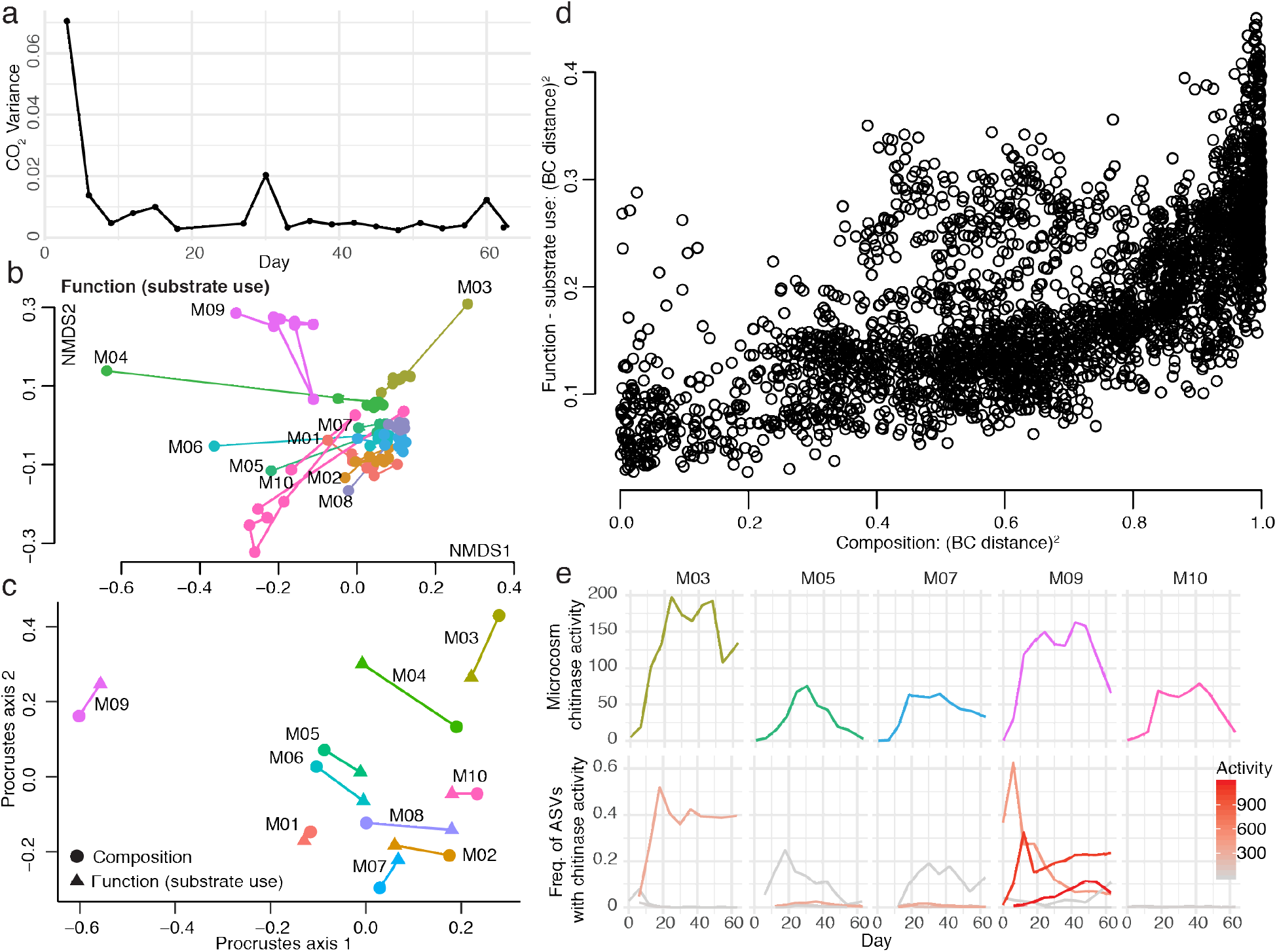
Community composition strongly correlates with functional activity. a) Variance in percent CO_2_ among the ten microcosms over the course of the serial transfer experiment. b) NMDS plot of the Bray-Curtis dissimilarities of functional activity (substrate use) as measured by EcoPlates. c) Procrustes rotation of the composition NMDS plot with the function NMDS plot for Day 63 (Procrustes correlation = 0.8991, p < 0.001). d) Plot of Bray-Curtis dissimilarities for composition by function (Mantel test r = 0.6403, p < 0.001), squared to better illustrate the spread of points. e) Endochitinase activity over time for the five microcosms that strains were cultured from (top row) and frequency of the ASVs over time that map to strains with measurable endochitinase activity (bottom row). Endochitinase activity is shown in units/mL for the strains/ASVs in the bottom row by a gradient from gray (low activity) to red (high activity).

In contrast with CO_2_ production and the expectation of functional convergence, the stabilized microcosm communities showed clear differences in substrate use, as measured across 31 substrates with Biolog EcoPlates (Supplementary Figure S3, Figure 4b and 4c). For example, M09 was the only community able to metabolize 2-Hydroxybenzoic acid (salicylic acid) but unable to metabolize itaconic acid. When plotting an NMDS ordination of Bray-Curtis dissimilarities based on EcoPlate functional measures (Figure 4b) we found a very similar pattern as with species composition: functional activity undergoes an initial large shift and becomes more similar, yet remains distinct across microcosms. A Procrustes test comparing the NMDS plots of composition and function at the end of the experiment recovers a strong and highly significant correlation: 0.8991, p < 0.001 (Figure 4c). Furthermore, when comparing composition to function across all days and samples using Bray-Curtis dissimilarities, samples with similar composition generally also had similar functional activity (Figure 4d), and are strongly correlated in a Mantel test (r = 0.6403, p < 0.001). The correlation is in fact higher when only comparing the final day’s measurements (r = 0.6907, p < 0.001). To profile the hydrolytic activity of the community, we focused on the activity of chitinases – the enzymes that degrade chitin – since chitin is the main component of insect exoskeletons and is a key carbon and nitrogen source in the pitcher plant system. The chitin hydrolysis rate of the community supernatant was also highly variable across microcosms, with M03 and M09 having the highest measures of endochitinase activity (Figure 4e). In summary, microcosm communities begin with different compositions as a result of historical contingencies affecting individual pitchers. These compositions shift as bacterial communities are brought into a new laboratory environment, but remain influenced by their starting compositions (Figure 5, part i). We expected to see functional convergence (ii), since all communities were grown for many generations in the same environment, but instead communities retained functional differences that were correlated with their compositions (iii).

**Figure 5.**
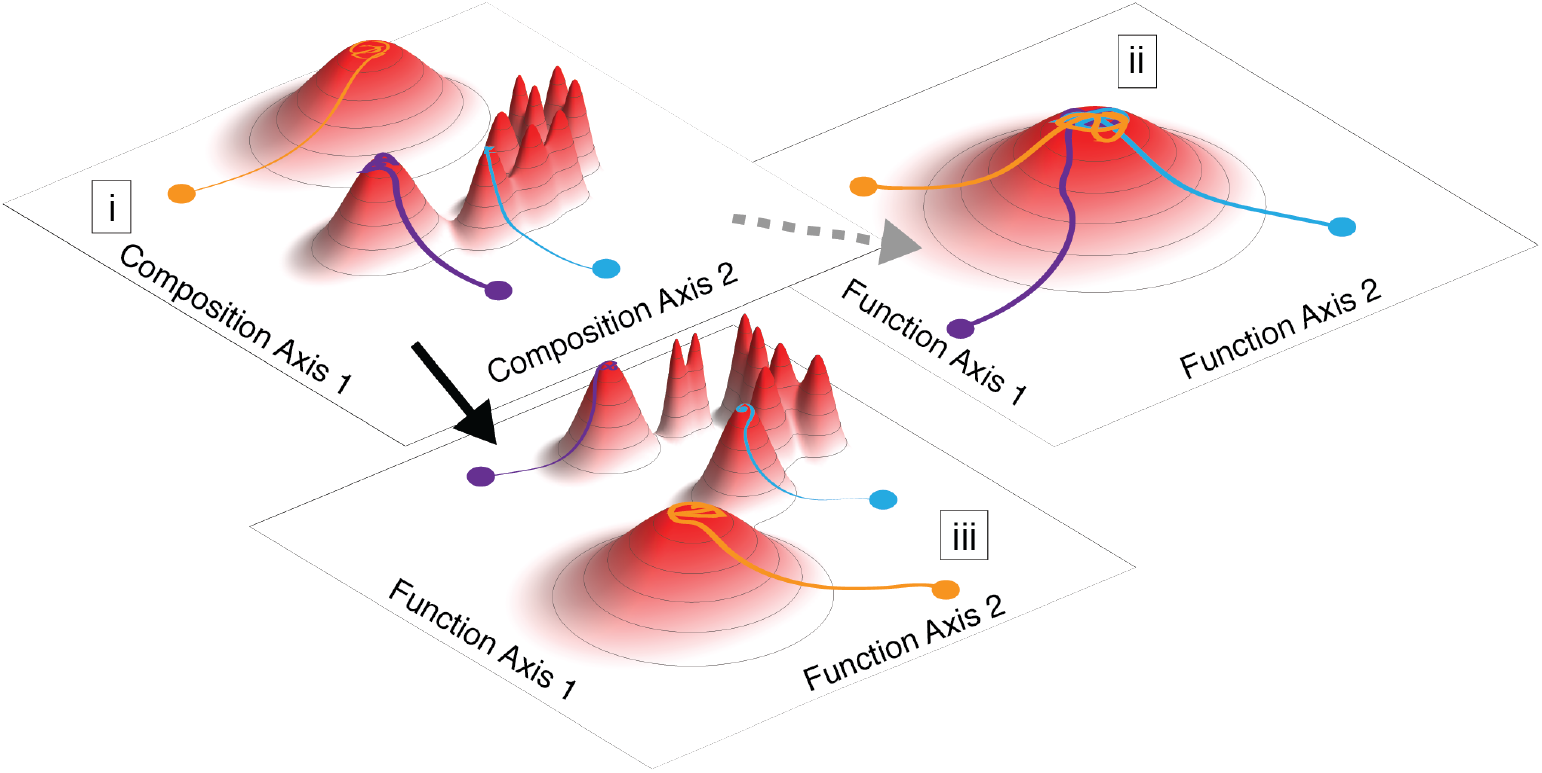
Microcosms with the same environmental conditions assemble communities with distinct equilibria due to historical contingencies (i). Function is often expected to converge in a common environment (ii), but our results show key functional differences are maintained by communities of different compositions (iii).

We developed an isolate collection to learn more about the mechanisms underpinning the strong differences in substrate utilization and hydrolytic activity across microcosms. By testing the enzymatic activity and substrate metabolism of individual isolates, we hypothesized we could identify bacterial strains responsible for the corresponding community function. We successfully isolated 350 strains and sequenced their full 16S rRNA genes. Of these, 176 mapped with 100% identity to 33 different ASVs from the amplicon sequencing. For the five microcosms we cultured from, 14 out of the combined top 15 ASVs (in terms of relative abundance) on Day 63 matched perfectly with cultured strains, as did 5-7 of the top 10 ASVs per microcosm. Our cultured strains had broad representation across different taxonomic groups (Supplementary file).

Consistent with our hypothesis, the chitinase activity of our cultured strains mirrored microcosm activity: strains from M03 and M09 had the highest endochitinase activity out of all measured strains. The activity of individual strains/ASVs mapped well to the activity of the entire microcosms, where M03 and M09 were also highest (Figure 4e). In M03 only one of the measured strains showed high activity (ASV589, a *Chromobacterium* species), while in M09 at least three strains had high endochitinase activity (ASV018 *Chromobacterium*, ASV842 *Dyella*, and ASV863 *Burkholderia*). Interestingly, the strains with the highest chitinase activity also had high protease activity, and all except for ASV842 had high lipase activity (Supplementary Figure S4), suggesting that these bacteria are generally good at degrading complex substrates.

To ask to what extent the pattern of substrate utilization of the community could be reduced to the substrate utilization of its members, we applied the same Biolog EcoPlates test to the individual isolates. We found that substrate utilization differences between communities could be attributed to differences in species abundance. Out of the 20 strains measured using Biolog EcoPlates, only one showed high metabolic activity when grown with salicylic acid (strain M09D5GC17 corresponding to ASV863 *Burkholderia*), and it was a strain present only in M09 which was the only microcosm where growth on salicylic acid was observed. Conversely, the strain with the most growth on itaconic acid (strain M10D5GC19 corresponding to ASV140 *Achromobacter*) was present at the final timepoint in all the microcosms we cultured from except for M09, and the M09 community was the only one not able to grow on itaconic acid (Supplementary Figure S5). Individual strains thus drive key functional differences among microcosms, and we were able to culture and analyze a set of these strains—connecting genotypes with their functional phenotypes.

## Discussion

The effects of historical contingencies are difficult to isolate and detect in the field, because even adjacent sites can experience distinct environmental conditions. The effects can be difficult to capture in a laboratory as well, because historical contingencies may only affect community assembly when disturbance is low and environmental selection is weak^2,12,35^ and these conditions are rarely met when natural communities are moved into laboratory settings. Our study demonstrates that historical contingency strongly influences community assembly in a realistic, but controlled, laboratory environment. Moreover, the effects of historical contingency are persistent and reproducible, suggesting that, with enough information about species’ functional capabilities, responses to environmental conditions, and interactions with other species, community assembly dynamics might be predictable.

We find the dynamics of individual ASVs affected key functional measures and were influenced by community context; likely driven by interactions among species within microcosms. For example, priority effects may have played a role, with early colonizers altering growth conditions for other species. Microcosm communities remained different in richness and in composition, despite an initial shift when assembling in a constant *in vitro* environment. Surprisingly, strong differences also remained in terms of relevant functional activity and function was highly correlated with composition in our study. These results suggest that species across the microcosm communities were not functionally redundant with regard to the relevant substrate degradation capabilities of this system. Redundancy in bacterial functional roles has been suggested as an explanation for the high complexity of microbial communities, and as a buffer that increases stability in the face of perturbation^34,36^. This study highlights how specific and relevant functional measures should be examined more closely in microbial ecology, because a general lack of redundancy in key functions could influence both the carbon flux and the overall stability of an ecosystem.

Our conceptual model for the assembly dynamics of microcosms suggests that when in the same *in vitro* environment, non-metabolically active species are quickly pruned, and after only one transfer the final diversity of the community is determined. This result implies that historical contingencies influence richness levels, even after communities equilibrate to a common environment. Despite differences in richness and composition across microcosms, temporal dynamics of ASV extinction follow the same processes, likely driven by external factors in the transfer process rather than the biotic context. However, ASVs persisting within the microcosms are largely influenced by microcosm context, indicating significant effects of species interactions. Therefore, long-lasting effects of early conditions and biota lead to strong differences in final community composition and ecosystem function. The environmental conditions of our experiment supported multiple functional outcomes, which may have shifted the selective balance to species interactions, therefore increasing the possible community states. Our experiment was necessarily run in the laboratory, but it used wild communities as the starting point and suggests potential implications for a natural system. Stochastic events during colonization of the pitchers of carnivorous pitcher plants may have lasting impacts on the ability of the pitcher microbiome to degrade insect prey and to release nutrients such as nitrogen and phosphorus to the common pitcher pool in a plant-accessible form.

Our model system based on pitcher plant bacterial communities can be used to address other questions in microbial ecology. For example: the role of dispersal in community assembly; how coalescence events (the mixing of stable communities) lead to different compositions; how invasions alter community structure and function; and how evolution acts on individuals within communities to change interactions over time. Our microcosms are less complex than most natural systems because species that did not grow within the current environment were pruned during transfers, but are more complex than almost all experimental laboratory communities. The ability to culture key community members provides the opportunity for building synthetic communities that retain interactions among species previously established in nature.

## Methods

### Sample collection

We collected the entire aquatic pools from 10 healthy pitchers of *Sarracenia purpurea* pitcher plants at Harvard Pond (Harvard Forest, MA) in September, 2017. We used sterile, single-use pipettes to remove the samples into sterile 15 mL tubes. The samples were transported in a cooler on ice to the laboratory where they were refrigerated overnight. The following morning, we set up the experiment.

### Serial transfer experiment

We filtered half of each sample through 3 μm syringe filters to focus on the bacterial component of the community. From both the filtered and unfiltered components of each sample, we combined 500 μL of pitcher fluid with 500 μL of media in a 48-well plate. In order to have a complex nutrient source similar to what bacteria from pitcher plant fluids would experience in their native environment, we used cricket media (3 grams cricket powder from farmed *Acheta domestica* crickets per 1 Liter of milliQ-purified water, acidified to pH 5.6 and then autoclaved). The plate was then placed in a 25°C incubator. After three days of incubation, each sample was mixed well and 500 μL was transferred to a new plate with 500 μL of sterile cricket media. From our calculations, we added the equivalent of about 1/60^th^ of a cricket to every well at each transfer. We continued transferring samples and adding cricket media in a 1:1 ratio every three days for a total of 21 plates over 63 days.

At the beginning of the experiment (Day 0), we removed a portion of each sample to freeze at −80°C for later DNA extraction and amplicon sequencing. We removed 100 μL of each filtered sample to first measure optical density (OD) at 600 nm, and then used a Fluorimetric Chitinase Assay Kit (Sigma-Aldrich) to measure the activity of three different types of chitinases: endochitinases, chitobiosidases and β-N-acetylglucosaminidases. We bead-beat each sample for 1 minute and centrifuged it before using a portion of the supernatant in the assay, with two replicates for each sample. Our downstream analyses focused on endochitinases, the enzymes that cleave intramolecular bonds forming new chain ends.

We also measured a “functional fingerprint” of the communities with Biolog EcoPlates^37^. We diluted each sample 1:10, combining 1 mL of each filtered sample with 9 mL of purified water that had been acidified to pH 5.6 and then autoclaved. We filled each EcoPlate well with 100 uL of sample, and incubated the plates in the 25°C incubator for three days. At the end of this time the plate was read in a plate reader according to EcoPlate instructions.

In addition to measuring chitinase and EcoPlate activity, we measured CO_2_ production with the MicroResp system^38^. We added 250 μL of each filtered sample to 250 μL of cricket media in three replicates for each sample in a deepwell plate, attached a detector plate with the MicroResp seal and clamp, and incubated at 25°C for three days before measuring the resulting color change in the detector plate at 570 nm. We calibrated the MicroResp measurements according to the manual by incubating the detector medium with known CO_2_ concentrations and making a reference curve.

At each transfer, we repeated the MicroResp measurement and froze a portion of the culture at −80°C for later DNA extraction. Every second transfer, we repeated the chitinase activity measurements, and every third transfer we repeated the Biolog EcoPlates with a 1:40 dilution to reduce carry over of any remaining cricket medium. All measurements after the first sets were done without replicates. During the course of the experiment, some of the MicroResp indicator plates showed evidence of fungal contamination; measurements involving affected wells were removed from our analyses.

On the final day (Day 63) of the experiment, we repeated all functional measurements, cultured five of the ten microcosm communities in order to isolate strains, and froze 100 μL of each community in 40% glycerol solution and the remaining culture at −80 °C.

### DNA extraction and sequencing

DNA was extracted from all samples with the Agencourt DNAdvance kit (Beckman Coulter) using 100 μL per sample. In each 96-well extraction plate we included negative controls. DNA was quantified with the Quant-iT PicoGreen dsDNA Assay kit (Invitrogen) on a plate reader, before being sent to the Environmental Sequencing Facility at Argonne National Laboratory for amplicon sequencing on a MiSeq targeting the V4 region of 16S rRNA using the same 515F and 806R primers as the Earth Microbiome Project (http://press.igsb.anl.gov/earthmicrobiome/protocols-and-standards/16s/). Sequence data has been deposited in the NCBI Sequence Read Archive (SRA) under Project ID PRJNA559886.

### Amplicon sequence analysis

On the MIT Engaging computing cluster, we used QIIME2 version 2018.4^39^ to demultiplex our sequences, and the DADA2^40^ plugin to denoise sequences and to generate Amplicon Sequence Variants (ASVs) of ~250 base pairs in length. We retained all ASVs with more than two sequences across all samples. We assigned taxonomy using the classify-sklearn method which is a Naive Bayes classifier, and a pre-trained classifier made with the Greengenes database, version 13_8.

Statistical analyses were performed and graphs were generated in R and Mathematica. Reads for each ASV were normalized by the total amount of reads in each sample. Bray-Curtis dissimilarities and NMDS ordinations for Figure 1 were performed using the R *vegan* package^41^. The effective number of species was calculated as exp(Shannon index). Richness for Figure 2 was calculated counting all ASVs present at later time points in each microcosm as present at previous time points to account for ASVs below the sequencing detection limit. Richness of early timepoints (Days 0 and 3) were correlated with that of the final timepoint (Day 63) using linear regression. We used additional R packages, including *plyr*, *ggplot2*, *reshape2*, *dendextend*, and *biclust*. R code and data tables are available via the Harvard Dataverse: https://dataverse.harvard.edu/privateurl.xhtml?token=69114952-e9e3-469d-844d-7e3c70380cd0.

### Null model for community assembly

To describe the dynamics of ASV loss, we employed a geometric model, i.e., the probability *P*(*t*) of an ASV that goes extinct eventually to go extinct at transfer *t* is equal to the probability that it did not go extinct in the preceding *t* − 1 transfer. That is,

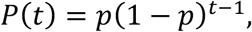

Where *p* is the sole parameter of the model. It can be shown that the maximum likelihood estimator of *p* is given by the mean time to extinction, i.e., 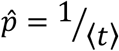. To determine whether the extinction time distributions are best described by a single parameter *p* or whether individual parameters for each microcosm are needed, we computed the Akaike Information Criterion 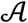 from the likelihood 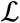 of the observed extinction under the geometric model, using either a single parameter *p* given by the inverse of the mean extinction time for all ASVs across all microcosms, or for each microcosm individually, i.e.,

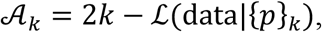

where *k* is either 1 (for a common parameter) or 10 (for individual parameters). We found 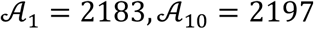, such that the relative likelihood of the common-parameter model over the individual-parameter model was 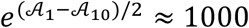.

### Correlation analysis

For the correlation analysis in Fig. 3c-e, ASV abundances were first center-log transformed after removal of absent ASVs. One pseudo-read was added to account for ASV abundances below the detection threshold. Standard Pearson correlation coefficients were then computed using the transformed time series. For Fig. 3c, we computed the correlation coefficients of time series of the same ASV in different microcosms and compared them to correlations between randomly chosen ASVs in different microcosms. For Supplementary Fig. correlation between pairs of strains, we computed correlations between pairs of strains that co-occurred in more than one microcosm and compared to randomly chosen pairs.

### Null model for an ASV having the same fate in multiple microcosms

For Fig. 3e, we classified the fate (either extinction of persistence) of individual ASVs shared between microcosms. We found that *f* = 80% of strains occurring in two microcosms had the same fate in both, suggesting that a given strain has an 80% chance of having the same fate in a new microcosm as it had in its current microcosm. We thus estimate the probability *P*(*n*) of a similar strain occurring in *n* microcosms to have to same fate in all *n* microcosms as *P*(*n*) = *f*^*n*−1^, which is shown as the red line in Fig. 3e.

### Strain isolation and identification

Individual strains were isolated from five of the ten microcosms (M03, M05, M07, M09 and M10) by plating the culture fluid and picking around 100 colonies per microcosm. We cultured on petri plates at 1:10^5^ and 1:10^6^ dilutions using both cricket media with the addition of agar and a second medium containing 11.28 g/L M9 salts, vitamin solution^42^, trace metals^43^, agar, and 2.5 g/L N-acetylglucosamine (GlcNAc) as the sole carbon source. Plates were put in a 25°C incubator for one week, after which single colonies were picked and then re-streaked at least two additional times before being grown up in liquid media and frozen in glycerol at −80°C.

To identify and barcode our strains, we added 2 μL of liquid culture to 20 μL of sterile, nuclease free water and, after a freeze-thaw cycle, did direct PCR using primers 27F and 1492R to amplify the 16S ribosomal RNA gene and the Q5 High-Fidelity kit (New England Biolabs) with an initial 5-minute incubation at 100°C. Before Sanger sequencing, we tested for successful amplification with gel electrophoresis and cleaned the PCR products with SPRI beads according to the Agencourt AMPure XP protocol. Sanger sequences were trimmed and filtered with Geneious, and assigned taxonomy using the RDP classifier.

### Measurements of strain functional activity

We measured the functional activity of approximately 50 strains with 100% matches in their Sanger-sequenced 16S rRNA gene to ASVs from the amplicon sequencing. When multiple strains mapped to the same ASV, their enzyme activities were averaged. Each strain was streaked out on cricket-M9 media plates from the frozen glycerol stock, and then a single colony was grown in liquid cricket media. Chitinase activity was measured as described for the microcosm communities, and for Figure 4e strains were considered to be active above a cutoff of 1 unit/mL. Protease and lipase activities were also measured using the Sigma-Aldrich Protease Fluorescent Detection Kit and Lipase Activity Assay Kit III. Seventeen of the strains were put into EcoPlates to compare their metabolic activity on the 31 substrates to that of their source communities.

## Supporting information

Supplementary Table of Strains

## Acknowledgements

We thank Jose Saavedra for help with sampling, Lei Ma and Peter Duff for laboratory assistance, and Veda Khadka for contributing R code. This work was made possible by NSF-BSF grant DEB 1655983 and European Research Council grant 640384 under the European Union’s Horizon 2020 research and innovation program. LSB was supported by a James S. McDonnell Foundation Postdoctoral Fellowship, MG was supported by the Simons Foundation Award 599207 and G.E.L. was supported by the Human Frontiers Science Program grant LT000643/2016-L.

