## Supplementary Figures for "Context-dependent dynamics lead to the assembly of functionally distinct pitcher-plant microbiomes"

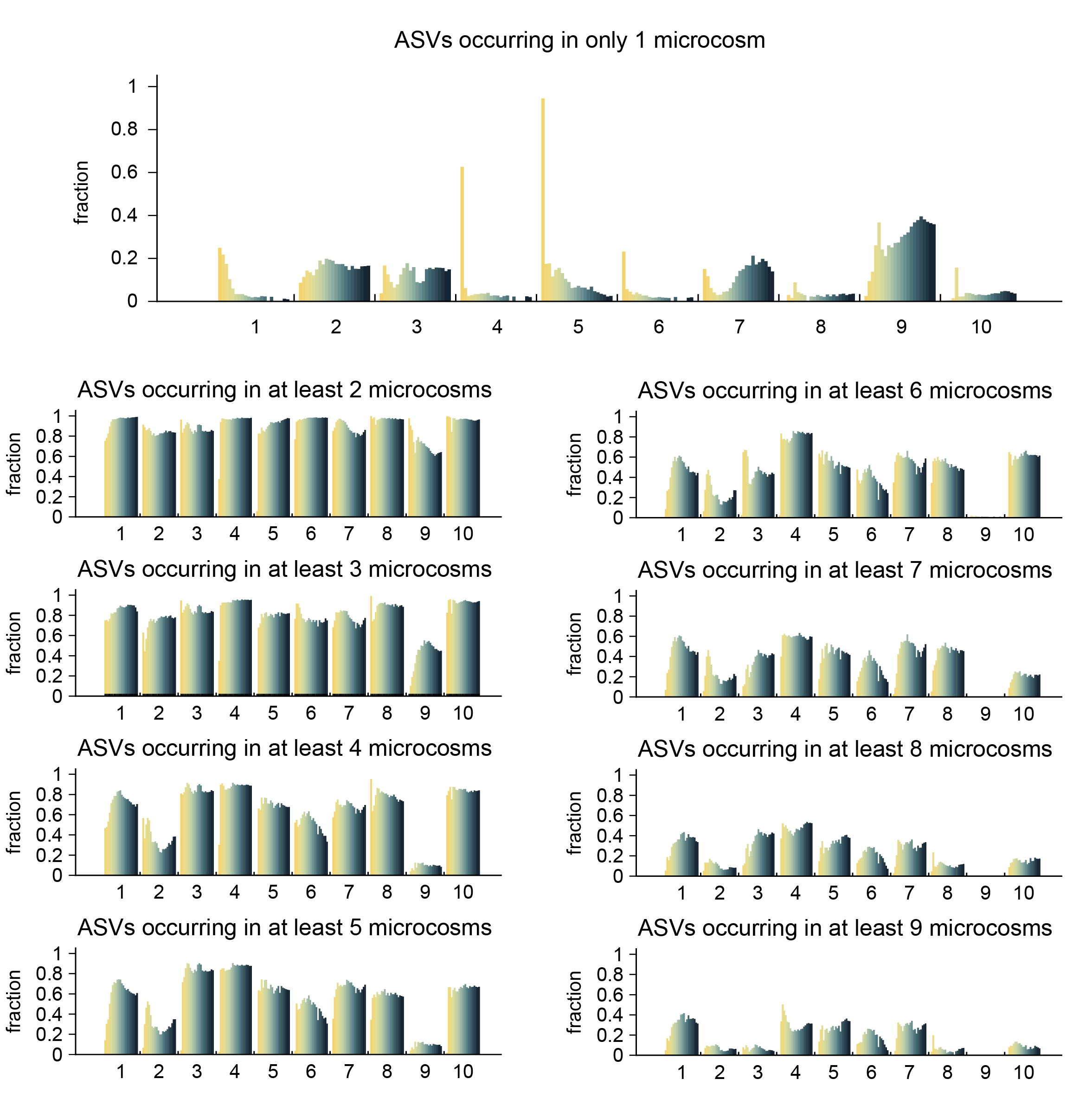

**Supplementary Figure S1**. The fraction of each microcosm contributed by ASVs occurring in *n* microcosms, shown over time (from Day 0 in yellow to Day 63 in dark green). M09 has the largest fraction of unique ASVs (~25%, top panel), while M01, M04, M05, and M07 are to a large degree (~40%) composed of ASVs occurring in all but 1 microcosm (bottom right panel).

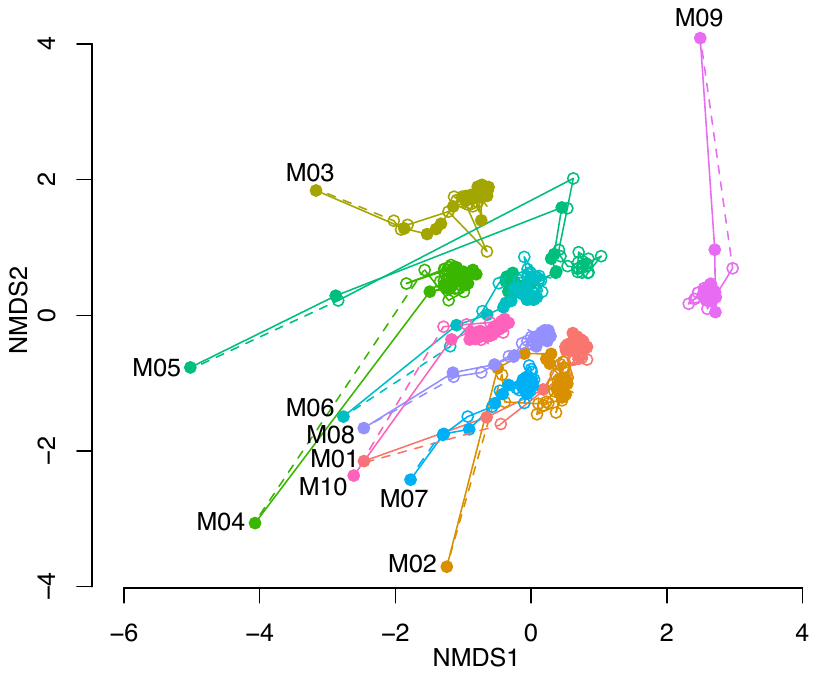

**Supplementary Figure S2**. Effects of environmental selection and historical contingency are replicated in both filtered and unfiltered microcosm communities. Filtered (solid circles) and unfiltered (open circles) samples follow almost the same trajectories, and are very similar in the NMDS ordination of their Bray-Curtis dissimilarities over time. The dashed line represents a suggested origin, as unfiltered communities were not sequenced for Day 0.

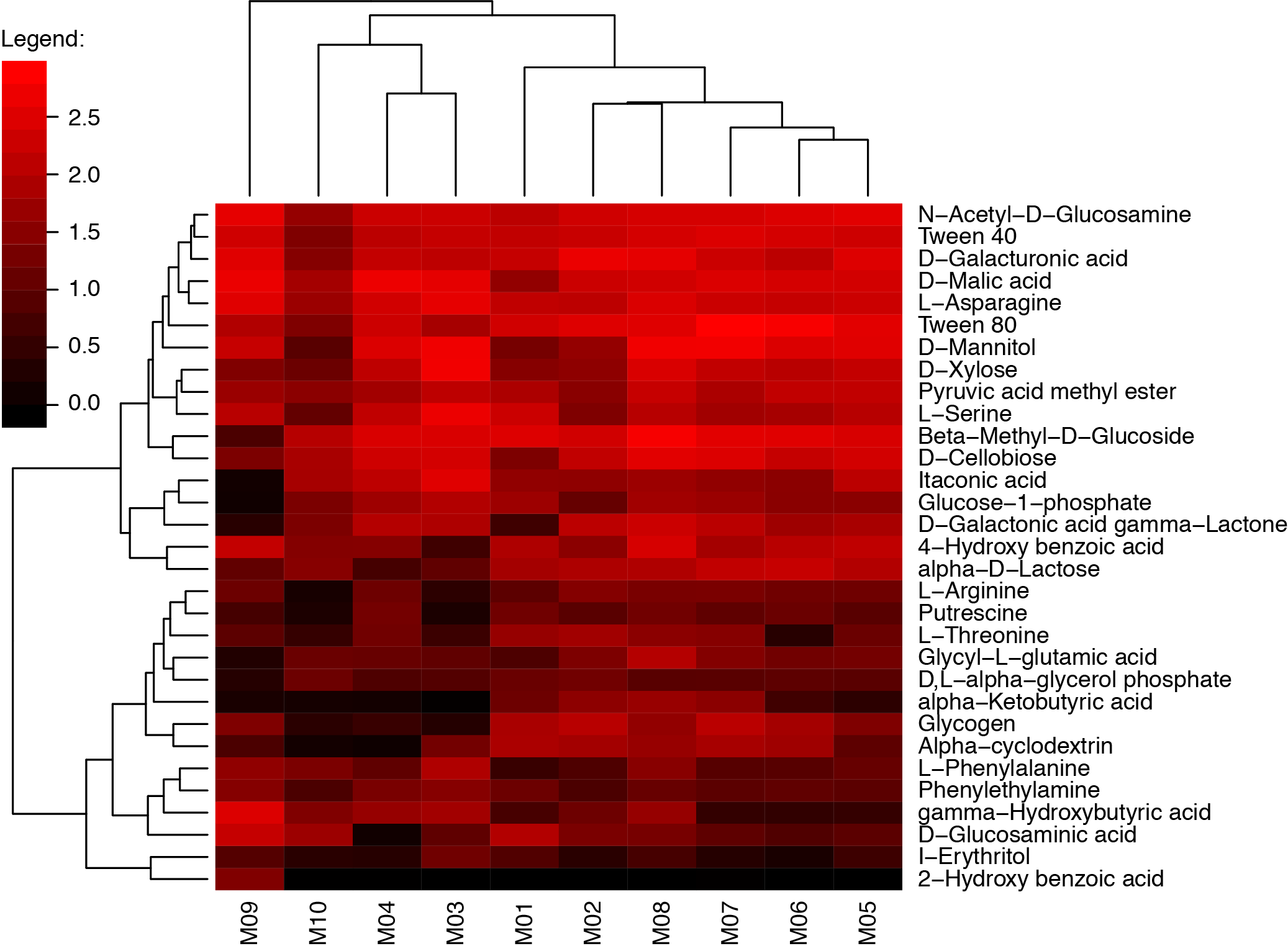

**Supplementary Figure S3**. Heatmap of EcoPlate data for stabilized microcosms (averaged across Day 27 to Day 63). Microcosms (columns) and substrates (rows) are arranged by similarity according to hierarchical clustering.

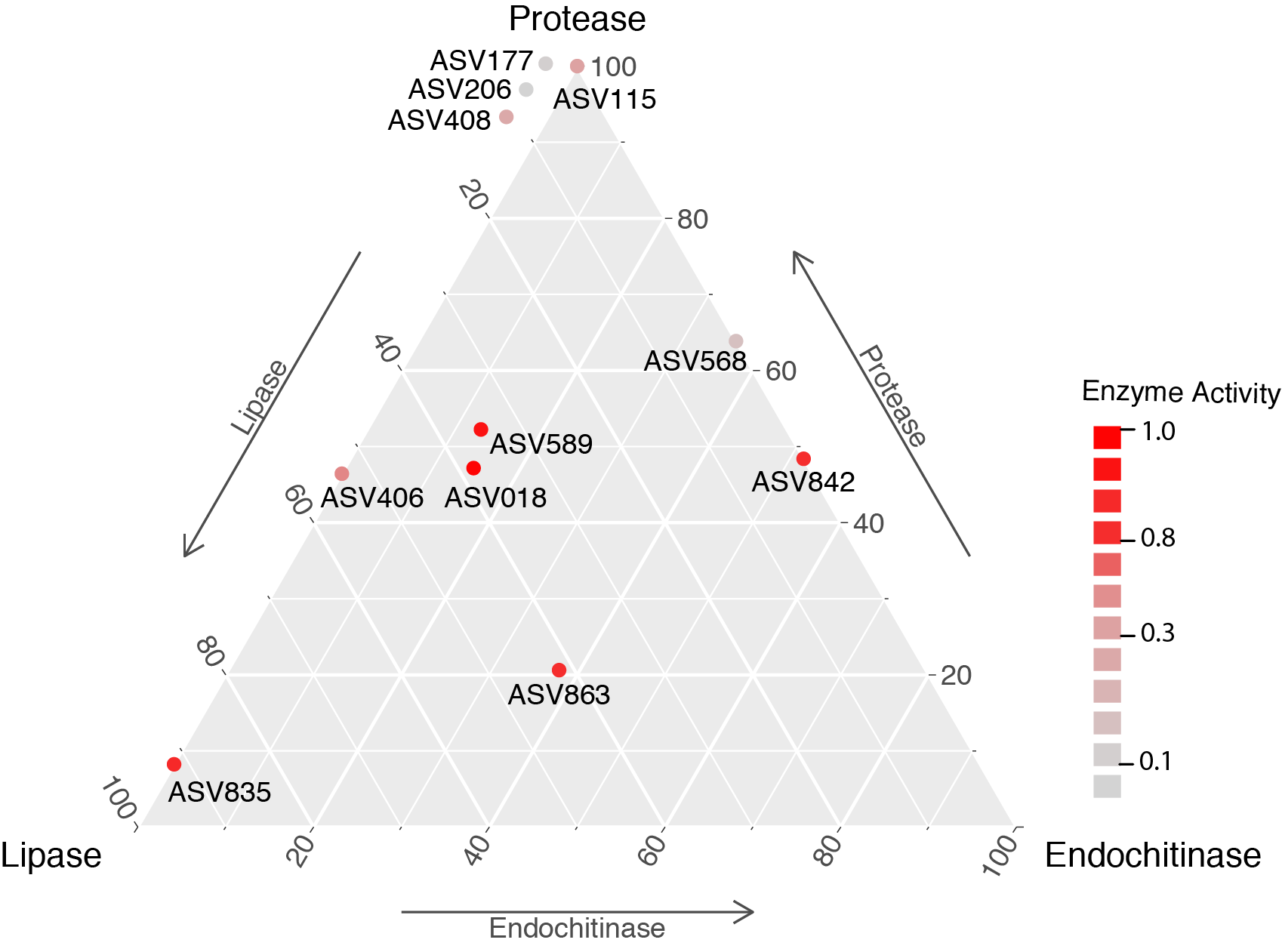

**Supplementary Figure S4**. Ternary plot of enzyme activity (endochitinase, lipase and protease) in cultured strains, mapped to corresponding ASV. Enzyme activity was normalized across strains and only values about 0.04 are included in this plot. Strains with high chitinase activity generally also have high protease and lipase activity (positioned near the center of the plot).

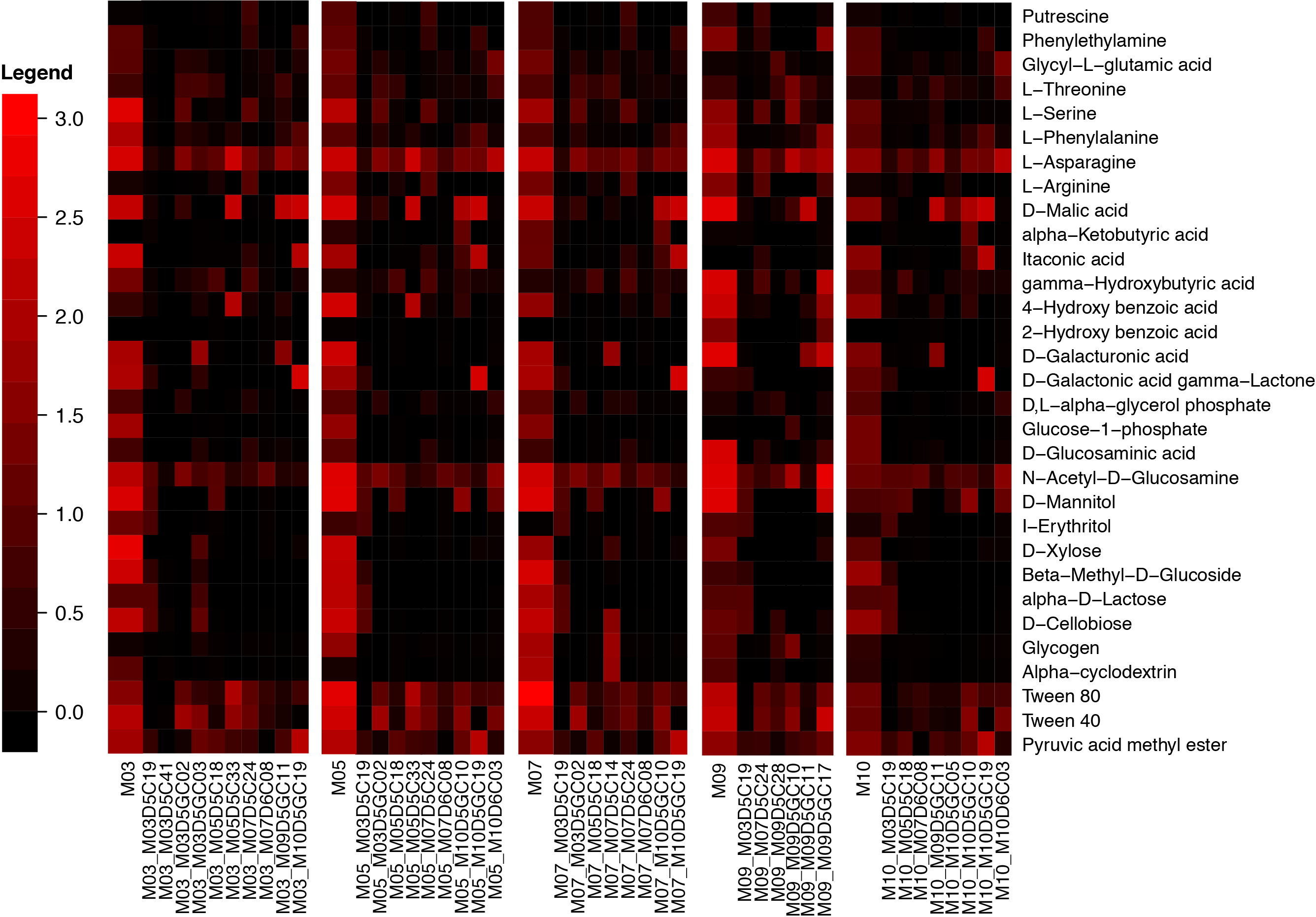

**Supplementary Figure S5**. Heatmap of EcoPlate results for Day 63 of Microcosms where strains were cultured, as well as a subset of the cultured strains (organized by which were present in each microcosm). Rows show EcoPlate substrates, columns are either communities or strains. The legend shows how color corresponds to substrate use: black is substrate not used, red is high use.
